# Molecular analysis of Lancefield group C/G streptococci causing human infections in Sheffield, UK

**DOI:** 10.64898/2026.01.30.702767

**Authors:** Saikou Y Bah, Henna Khalid, Sona Jabang, Roy Chaudhuri, Lisa Tilley, Luke R Green, David Patridge, Thushan I de Silva, Claire E Turner

## Abstract

Lancefield group C/G streptococci (GCS/GGS) are increasingly recognised as significant human pathogens that cause a disease spectrum similar to *Streptococcus pyogenes*. Despite their high clinical burden in the UK, their genomic diversity remains poorly understood. We performed whole-genome sequencing (WGS) on a prospective collection of 109 consecutive GCS/GGS isolates from all infection types in Sheffield, UK, over five months in 2020. *Streptococcus dysgalactiae* subsp. *equisimilis* (SDSE) accounted for 104 isolates, while five were identified as *Streptococcus canis*.

The SDSE population was highly diverse, comprising 15 genomic clusters and 38 unique *emm*-ST combinations. We identified the presence of the ST20/stG62647 international lineage (24% of isolates), a cluster globally associated with severe invasive disease. Antimicrobial resistance genes were prevalent (49%), predominantly linked to mobile genetic elements carrying tetracycline and macrolide resistance. Furthermore, a variation in the penicillin-binding protein PBP2X (P601L) was linked to reduced penicillin sensitivity (MIC 0.03 mg/L). There were few or no genetic changes in isolates obtained from the same patient, even when they were collected 8-10 weeks apart, indicating long-term persistence within a host. The unexpected detection of *S. canis* in human infections and the high diversity of SDSE, persistence and virulence-associated regions underscore the need for enhanced national genomic surveillance to track emerging virulent and antibiotic-resistant SDSE lineages.

**Impact statement:** Lancefield group C and G streptococci, most often the species *Streptococcus dysgalactiae* subsp. *equisimilis* (SDSE), are an increasingly significant human pathogen, often mirroring the severity of infections caused by Lancefield group A *Streptococcus* (*S. pyogenes*). Despite its clinical importance, we know little about the population of SDSE circulating in the UK. This study provides the first comprehensive genomic analysis of SDSE isolates from a single UK region, identifying a highly diverse population comprising 15 distinct genomic clusters but with evidence of long-term persistence within a single host. Notably, we confirm the presence of the international *stG*62647/ST20 lineage in the UK, which is globally associated with severe invasive disease. Our findings also reveal a high prevalence of antimicrobial resistance genes (∼49%), primarily linked to mobile genetic elements, and the presence of a specific variation in the penicillin-binding protein PBP2X that reduces penicillin sensitivity. Additionally, the unexpected detection of *S. canis* in human infections rather than animals highlights a need for monitoring. By defining the UK’s SDSE population structure and its resistance landscape, this research underscores the critical need for enhanced national genomic surveillance to track emerging high-virulence and antibiotic-resistant lineages

**Data summary:** Sequence files for isolates from Sheffield used for this study have been uploaded to the sequence read archive with project accession number PRJNA1333937 and accession numbers provided in Supplementary Dataset 1. The completed genome for SDE096 been deposited on GenBank with the accession number JBSXMJ000000000.

## Introduction

*Streptococcus dysgalactiae* subsp *equisimilis* (SDSE) is a beta-haemolytic streptococci that is closely related to the human pathogen *S. pyogenes*, sharing many virulence factors and mobile genetic elements (MGEs) with this species (1). Although historically described to be the cause of infections in animals, over the past decade increased incidence of SDSE in humans has been documented in many parts of the world and in some geographical regions even overtaking *S. pyogenes* as a cause of invasive diseases (2, 3). Like *S. pyogenes*, SDSE can cause a wide range of infections in humans, including mild and self-limiting throat, skin and soft-tissue infections but also much more severe and potentially lethal invasive infections such as bacteraemia, necrotising fasciitis and toxic shock syndrome (1, 4, 5).

SDSE can carry the Lancefield group carbohydrate C or G, but is now recognised to occasionally carry group A, confounding species identification using standard microbial methods, as group A was thought to be restricted to *S. pyogenes*. SDSE also carries an *emm* gene, encoding for the surface protein M. The hyper-variable region of the *emm* gene is used for *emm*-typing in the same manner as *S. pyogenes*, and to date more than one hundred *emm*-types have been identified for SDSE with few of these also identified in *S. pyogenes*, possibly due to homologous recombination (6, 7).

Although group C and group G streptococci combined cause almost as many cases of bacteraemia in England and Wales as group A *Streptococcus*, (8) nothing is known about the genetic diversity of SDSE in the UK. Recent data from other European countries and elsewhere have consistently identified a lineage with the *emm*-type *stG*62647 and multi-locus sequence type (ST) 20 as a predominant cause of disease, including severe and invasive diseases (6, 9-18). It is not currently known if this lineage circulating in the UK. To try and address the lack of knowledge regarding the SDSE population in the UK, we collected all Lancefield group C and G streptococci identified at the microbiology diagnostic laboratory at the Northern General Hospital, Sheffield, UK in the summer (June-October) of 2020. This hospital diagnostic laboratory services the community as well as all of the city’s hospitals. We performed comprehensive genetic characterisation of these 109 group G/C streptococci, confirming the presence of the *stG*62647/ST20 lineage as a major cause of infection in addition to other diverse lineages.

## Methods

### Isolates

All Lancefield group C/G streptococci that were routinely diagnosed from any infection type using Lancefield typing latex agglutination and MALDI-TOF by the department of Laboratory Medicine, Northern General Hospital, Sheffield, UK between June and October 2020 were stored (n=109). Laboratory Medicine is UKSA accredited and the single regional microbiology diagnostic laboratory for Sheffield NHS trusts as well as primary and community care, servicing around 600,000 adults and children. Isolates were fully anonymised, and no clinical or patient data was obtained apart from a general description of the original site of isolation (either skin, throat, blood, genital or other).

### Whole genome sequencing

Stored isolates were sub-cultured on Columbia horse blood agar (Oxoid) and then in Todd-Hewitt media (Oxoid) for DNA extraction. DNA was extracted using a previously described method (19) and sent to MicrobesNG (https://microbesng.com) for short read whole genome sequencing. Sequencing libraries were prepared using the Nextera-XT library preparation kit (Illumina) and sequenced using the NovaSeq 6000 platform generating 250-bp paired end reads. One isolate, SDE096, was subjected to Oxford Nanopore long read sequencing. DNA was prepared for sequencing using the Oxford Nanopore Technologies (ONT) ligation sequencing kit (SQK-LSK109), according to the manufacturer’s protocols. Libraries were sequenced with the ONT MinION Mk1C using rev C 9.4 flow cells and run for 24 hours. Resulting trace files were analysed with the ONT Guppy basecaller (version 3.2.10) using high accuracy basecalling. A complete single-contig genome was assembled by combining the long read sequence data with the short-read Illumina data using Unicycler (v0.5.1) (20). The completed genome was submitted to GenBank for annotation (accession number: JBSXMJ010000000).

### Bioinformatics analysis

Illumina reads were trimmed using Trimmomatic (v0.38) (21) with the settings: LEADING:3 TRAILING:3 SLIDINGWINDOW:4:15 MINLEN:36. Draft assemblies were generated using Unicycler (v0.5.1) (20) and assembly statistics checked with Quast (v5.3.0) (22). All assemblies passed the quality control (all fewer than 120 contigs and with an overall length of 2.0-2.3 Mb). Prokka (v1.14.6) (23) was used to annotate the draft assemblies and core genomes determined using Panaroo (v1.5.2) (24) with strict mode. Single nucleotide polymorphisms (SNPs) to infer genetic distances between assemblies were determined from the Panaroo-generated core genome alignment using snp-dists (v0.7.0; https://github.com/tseemann/snp-dists). The sequences in the pan_genome_reference.fa generated by Panaroo were submitted to EggNOG (v2) (25) for functional annotation and grouped into cellular process, metabolism, information processing or uncharacterised. This output was visualised with Sankey plot generated with python plotly. Maximum likelihood phylogenetic trees were generated using a general time-reversal (GTR) substitution model with 100 bootstraps using RAxML (8.2.12) (26) and then visualised using iTol (v7.2.1) (27).

### Typing and cluster analysis

The *emm*-types were determined using the emm_typer.pl script available at https://github.com/BenJamesMetcalf/GAS_Scripts_Reference and the multi-locus sequence types (MLST) determined using the mlst_check from the Sanger institute available at https://github.com/sanger-pathogens/mlst_check. All novel alleles and STs were submitted to the pubMLST database (https://pubmlst.org/organisms/streptococcus-dysgalactiae). Population Partitioning Using Nucleotide K-mers (PopPUNK) (28) was used to cluster the SDSE draft assemblies using a reference database available at https://www.bacpop.org/poppunk-databases/. For further analysis of genome clusters, reads for the isolates within this cluster were mapped to a reference genome or the *de novo* assembly of one of the cluster members, and core SNPs were determined using Snippy (v4.6.0) (https://github.com/tseemann/snippy), and regions of recombination determined using Gubbins (3.1.0) (29) and visualised using Rscript plot_gubbins.R. Further genome sequence data for the *stG*62647/ST20 lineage was obtained from NCBI using the list of isolates identified to be within this lineage by Xie et. al 2025 (11).

### FCT Regions

To determine the FCT region, the location of some FCT genes were identified from the annotated *de novo* assemblies and the surrounding full FCT region was then extracted. Extracted regions were subjected to BLAST search of the entire NCBI database to identify all the genes within the region and then the order of these genes was used to assign FCT gene arrangement pattern types compared with previously determined FCT regions (1). Some samples had contig breaks in the extracted region and therefore a pattern could not be determined.

### Virulence factor, regulator and antimicrobial resistance typing

The presence of 13 *S. pyogenes* superantigen genes (30), the SDSE version of *speG* (*speG*_*dys*_) gene (31), and the DNase genes *sda1, sda2, sdn, spd1, spd3* and *spd4*, were determined by BLAST (100% coverage, 80% identity) against *de novo* assemblies with representative gene sequences and manual checks where necessary. The CovR/S genes were extracted from the draft assemblies using *in silico* PCR (https://github.com/simonrharris/in_silico_pcr), translated to amino acids and an in-house python script was used to identify variants. Antimicrobial resistance genes were identified using ABRicate (https://github.com/tseemann/abricate) using the NCBI AMRFinder plus database (32). The penicillin binding protein genes encoding for PBP1A, PBP1B, PBP2A and PBP2X were extracted from the *de novo* assemblies and compared as for CovR/S.

### Penicillin susceptibility testing

SDSE isolates were grown overnight in Todd-Hewitt broth (THB, Oxoid) at 37°C with 5% CO_2_. Cultures were adjusted to OD=0.1 in fresh THB, before diluting further 1/100 in THB. 75 µl of cultures were added to the wells of a 96-well plate (Corning), along with 75 µl of penicillin G (Merck) equating to final antibiotic concentrations ranging from 0.004 – 0.05 mg/L in 0.001 mg/L increments up to 0.01 mg/L then 0.01 mg/L increments. Plates were incubated for 24 hours at 37°C with 5% CO_2_. The MIC was the lowest concentration of penicillin G that was found to inhibit the growth of each SDSE isolate.

## Results

### Diversity within the population

After quality control filtering, we had *de novo* assemblies for 109 whole genome sequenced Lancefield group C/G isolates from 101 participants. Although all obtained on different dates, up to 10 weeks apart, three isolates were from one patient, and two isolates were obtained from each of a further 6 patients. 104/109 isolates were determined to be SDSE, predominantly isolated from skin (n=60, 57.7%) or from genital swabs (n=20, 19.2%) with low numbers of blood isolates (n=7, 6.7%) or throat isolates (n=4, 3.8%) (Supplementary dataset 1). Unexpectedly, five isolates were determined to be *S. canis*, a species thought to be a rare cause of disease in humans (33), isolated from skin (n=2) or other sites (n=3) and all from different individuals.

Pangenome analysis of the 104 SDSE genomes identified 4,172 genes in total, with a core genome size of 1,602 genes (Figure 1). Orthologous grouping indicated that 47.1% of accessory genes were uncharacterised compared to 23.5% of the core genome. A large proportion of the core genome (38.7%) was predicted to be involved in metabolism, compared to just 11.0% of the accessory genes, but an almost equal proportion of core and accessory genes (14.9% and 14.5% respectively) were predicted to be involved in cellular processes (Supplementary figure 1).

**Figure 1.**
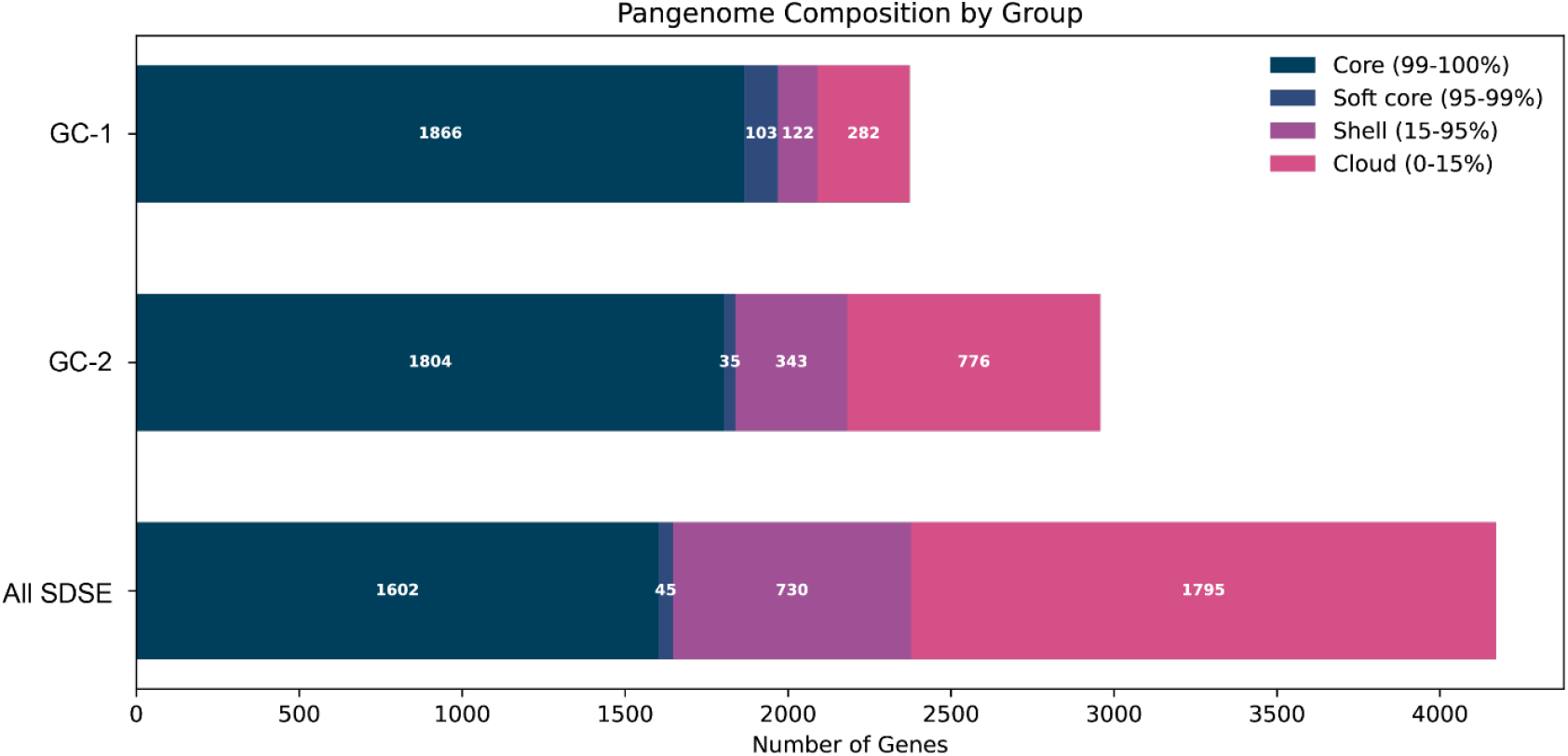
Pangenome analysis. Core and accessory genes were determined by Panaroo for all 104 isolates as well as the genomic cluster 1 (GC-1) and genomic cluster 2 (GC-2) isolates. The total number of genes for all the SDSE isolates, GC-1 and GC-2 were 4172, 2373, 2994 genes respectively.

Twenty different *emm*-types were identified in the 104 SDSE isolates; one isolate was deemed ‘non-typable’ due to a deletion in and around the *emm* gene locus. Those isolates obtained from the same patient were the same *emm*-type and MLST so therefore only a single isolate was used as representative per patient for further analysis. The *emm*-type *stG*62647 was the most common, accounting for 24.0% (n = 23/96) of the isolates, followed by *stC*74A, accounting for 18.8% (n = 18/96) of the isolates. MLST was also determined for the isolates, and twenty-seven different STs were identified. ST17 was the most common (34.4%) and eight different *emm*-types were found with this ST. ST20 was the second most prevalent type, accounting for 24.0% of the isolates but only one *emm*-type was associated with this ST (*stG*62647). Overall, there were 38 different *emm*-MLST combinations. Core gene phylogeny identified that, while the *stG*62647/ST20 isolates formed a single discrete lineage, multiple *emm*-types and multiple STs were found in other lineages (Figure 2). This finding is consistent with that of others (5), that *emm*-type and MLST are not good markers for the population structure of SDSE. We therefore additionally assigned genomic clusters to the population using PopPUNK. This produced fifteen genomic clusters, eight of which consisted of just one isolate. For the 12 *emm*-types that were found in two or more isolates, only 4 (*stG*10, *stG*839 and *stG*5345) were uniquely assigned to a single cluster, confirming the diversity within *emm*-types. There was no association between infection type and genome cluster but two genome clusters (1 and 2) were the most dominant.

**Figure 2:**
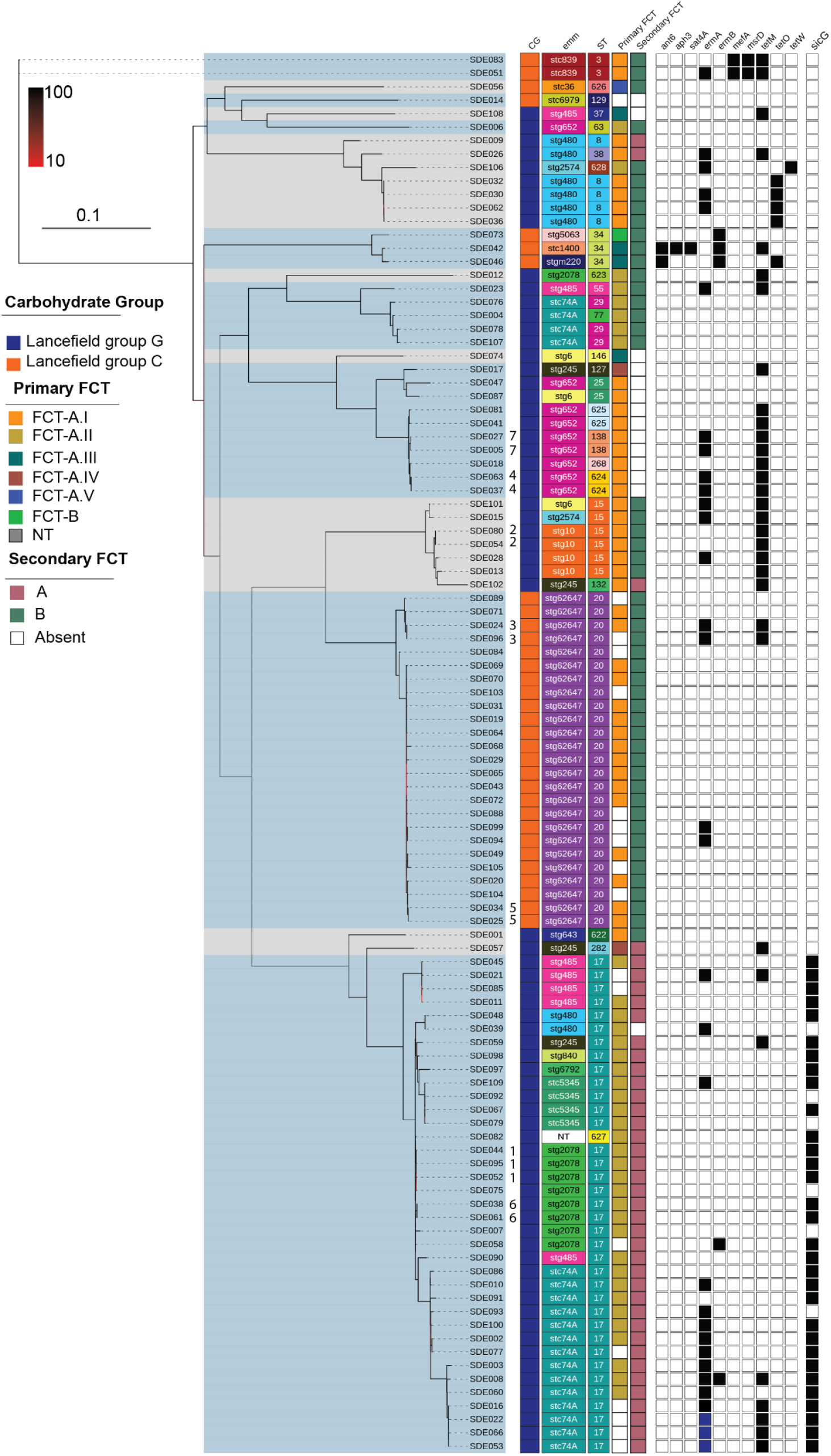
Phylogeny of the 104 Sheffield SDSE isolates. A maximum likelihood phylogenetic tree was constructed from the core genome SNPs using RAxML with 100 bootstraps. While Lancefield grouping of C (orange) or G (blue) was associated with PopPUNK genomic clusters (alternating grey/blue shading), *emm* and sequence type (ST) overall were not. The different primary and secondary FCT types are indicated, as are the presence (black) of AMR associated genes and the gene *sicG*. Isolates from the same patient are indicated by numbers 1-7, each number representing a different patient. Three isolates had two copies of *ermA* (blue). NT: non-typable. The scale bar (0.1) represents the number of nucleotide substitutions per site. Bootstrap scale (10-100) for branch colours is also provided.

### Antimicrobial resistance gene and penicillin binding protein gene variations

Tetracycline resistance genes were the most prevalent AMR genes identified within SDSE with 28/96 isolates carrying *tetM* (29.2%), a further 5 carrying *tetO* and one additional isolate with *tetW* (Figure 2). Macrolide resistance genes *ermA* and *ermB* were found in 29 (30.2%) and 5 (5.2%) of isolates respectively. These genes were found across the population and not restricted to single genome clusters, although three isolates in the same genomic cluster had two copies of *ermA* (Figure 2). The genes *ant6, aph3* and *sat4A* were together only in one isolate, although an additional isolate had *ant6* alone, and *mefA* and *msrD* were found together in two isolates of the same genome cluster (Figure 2).

Variations within penicillin binding proteins have been previously identified in *S. pyogenes* that conferred reduced sensitivity to beta-lactam antibiotics (34-37). Therefore, we extracted and compared the penicillin binding protein sequences (PBP1A, PBP1B, PBP2A and PBP2X) to identify any amino acid variations, focusing on the transpeptidase domain (Supplementary Dataset 1).

A number of different variations were found within PBP1A, including a variable number of tri-amino acid repeats towards the end of the protein. However, none of the 11 different variants identified were within the transpeptidase domain (residues 335-632, pfam: PF00905) and most of the combinations of variations were associated with genome clusters. This was also the case for PBP1B, including a common pattern of 16-17 amino acid variants found in clusters of isolates, but none of these variants were within the transpeptidase domain (399-635, pfam: PF00905). However, one isolate, SDE056, did have a variation within this domain (I413V).

Within the transpeptidase domain of PBP2A (422-685, pfam: PF00905) we found a common variation of T488M in all but one of the *stG*62647/ST20 genome cluster isolates as well as 10 other isolates from other clusters. One of these *stG*62647/ST20 genome cluster isolates also had A592G, and another one (of a pair from the same patient) had T561I. All but two ST17-containing genome cluster isolates had both T464A and I481V. These variations were also found in other isolates (Supplementary Dataset 1).

PBP2X was by far the most conserved with only six variations: V90L, D262N, A397V, R424Q, Q462K and P601L; the latter four are within the transpeptidase domain (292-603, pfam: PF00905). P601L has been previously reported in *S. pyogenes* and is associated with a reduction in penicillin sensitivity (37). This variation was found in the pair of *stG62647*/ST20 isolates from the same patient. The Q462V variation was found in all but five of the 34 ST17/ST627 isolates (genome cluster 2). This change has not been previously reported, although the variant Q462H was not associated with a change in sensitivity to penicillin in *S. pyogenes* (37).

To determine if any of the variations we identified in any of the *pbp* genes altered the sensitivity to penicillin, we measured the MIC to penicillin G. All isolates had an MIC of 0.02 mg/L or lower, apart from two isolates that had an MIC of 0.03 mg/L; these two isolates, from the same patient, had the P601L variation in PBP2X suggesting this variation slightly reduces the penicillin sensitivity of SDSE.

### FCT Pattern

The fibronectin-collagen binding-T antigen (FCT) genomic region contains genes encoding for pilin structural proteins, pilus biosynthesis enzymes, and other adhesins. This region can be highly variable and has been implicated in determining tissue tropism for *S. pyogenes* (38). The arrangement of the genes within the FCT region determines the FCT type. While only one FCT region has been identified in *S. pyogenes*, two FCT regions have been found in SDSE isolates (1, 39, 40). In our SDSE isolates the primary FCT loci had a gene arrangement similar to *S. pyogenes* FCT, all starting with the transcriptional regulator *rofA* (Figure 3). Six different patterns were identified, with five variants (I-V) based on the previously defined FCT-A (1, 40) depending on the variable presence of a fibronectin binding protein gene (*gfbA*) downstream of the transcriptional regulator, sortase genes and the pilin adhesin gene *pilA*. All FCT-A variants carried the pilin linker gene *pilL*, the pilin backbone gene *pilB* and a second fibronectin binding protein (*fbpZ*). FCT-B was found in a single isolate and was previously defined by the lack of *fbpZ* and a different order of the *pil* genes (1, 40). The most common, found in the most number of isolates, were FCT-A.I and FCT-A.II, without and with *gfbA* respectively. Broadly, FCT types were associated with genome clusters (Figure 2).

**Figure 3:**
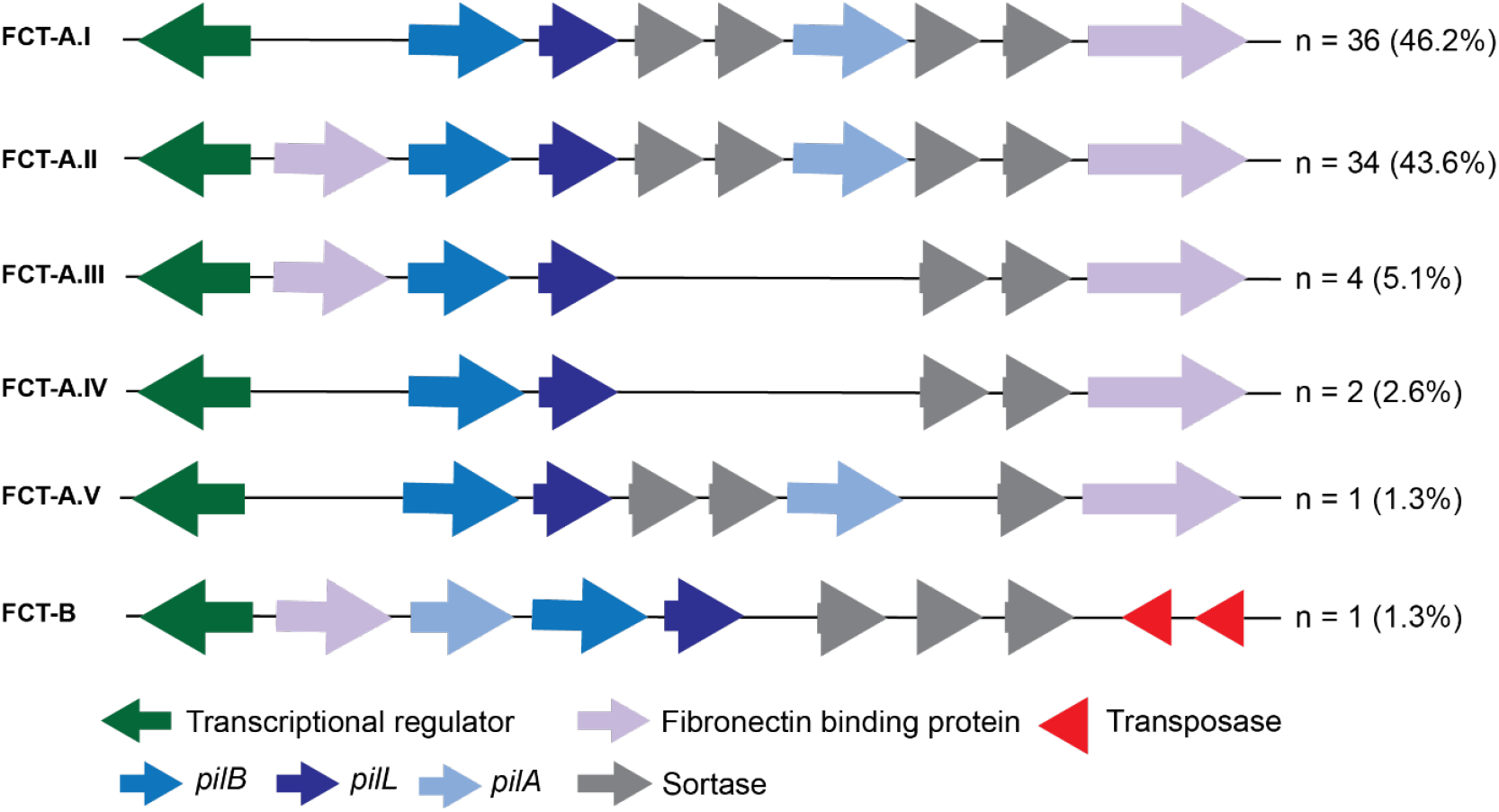
Gene arrangement of the primary FCT locus in SDSE. The FCT region was extracted within single contigs for 78/96 *de novo* genome assemblies. Five different variants of FCT-A (I -IV) and one variant of FCT-B were identified and the number of isolates (%) with these variants is indicated on the right.

A secondary FCT region was found in 90/104 isolates in two different variations depending on the presence of one copy of *srtB* and three pilin genes (variant A), or two *srtB* and four pilin genes (variant B) (Figure 4).

**Figure 4:**
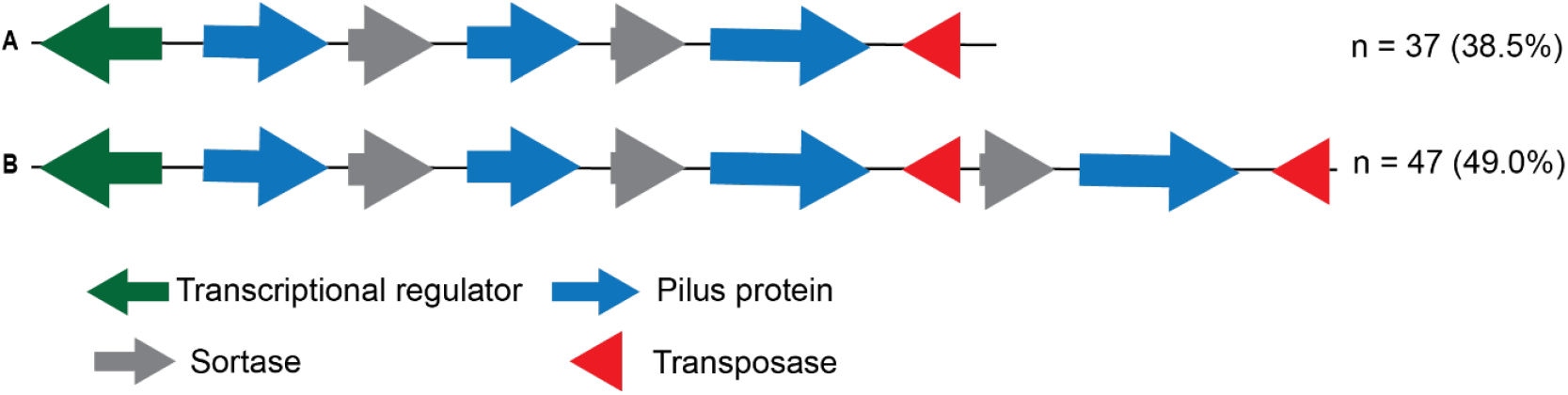
Gene arrangement of the secondary FCT locus in SDSE. Two variants of the secondary FCT region were identified in 84/96 isolates and the number of isolates (%) with these variants is indicated on the right.

### CovR/S

In *S. pyogenes* the CovR/S two component system controls the expression of ∼15% of genes in the genome and mutation in CovR or CovS are associated with hyper-virulence (41). A similar role for CovR/S has been identified in SDSE, repressing major virulence factors and potentially contributing to more severe disease (42-44). We found six isolates with unique variations in CovR: E9D, I50F, V88M, P103L, Y160F and S164G. These six isolates all belonged to five different genomic clusters suggesting they were spontaneous variations rather than lineage-associated and were either from skin infections (n=5) or other infections (n=1). Fourteen different amino acid change variations were identified in CovS in 17/96 (17.7%) of the isolate genomes, although combinations of some appeared to be associated with genomic clusters and we did not find any premature stop codons.

### DNases and Superantigens

*S. pyogenes* can carry a combination of 13 different superantigens (30), some associated with bacteriophages. The only superantigen we identified in our SDSE isolates was *speG*_*dys*_, which is the variant *speG* superantigen gene found in SDSE and differs from *speG* found in *S. pyogenes* (∼85% identity). It was present in 48 isolates, restricted to all isolates within ten genomic clusters. *S. pyogenes* can also variably carry prophage-associated DNases, and so we tested for the presence of *sda, spd1, spd3, spd4* and *sdn*. We found *sda, sdn* and *spd3* in three, one and three isolates respectively, although one isolate carried both *sda* and *spd3*.

### Diversity of the genome cluster 1 (*stG*62647/ST20)

Although it was the second most common cause of infections in our sampling, the genome cluster 1 (GC-1) exclusively consisting of ST20 and *stG*62647, has in other countries been a highly prevalent cause of infections, associated with severe disease, including streptococcal toxic shock syndrome (STSS) and necrotising soft tissue infections (6, 12, 13, 45). Here, we found the majority were associated with skin infections (13/25), including two isolate pairs from the same patients, all skin, and none were blood isolates.

Consistent with the conserved ST and *emm* within GC-1, there was a fairly low level of recombination (Supplementary Figure 2) and a low level of accessory genome content (Figure 1). They also all had, where we could determine the type, primary FCT-A.1V and the secondary FCT region variant B.

Comparison with other publicly available *stG*62647/ST20 isolates identified that our Sheffield *stG*62647/ST20 GC-1 isolates also belonged to this international lineage and were distributed throughout the tree, indicating no regional isolation (Figure 5). We sequenced one of our isolates, SDE096, with long read sequencing to complete the genome. As previously identified in *stG*62647/ST20 isolates, SDE096 has a transposon inserted into the *silB* gene, part of the streptococcus invasion locus (*sil*), mutations in which have been associated with more severe clinical outcomes (18). All other Sheffield GC-1 isolates also had this insertion in *silB*. The *sil* locus was found in 58/96 (60.4%) isolates in total but only GC-1 isolates had an insertion in *silB*, and other isolates had full length *silB*, apart from three which had premature stop codons truncating SilB after 82, 99 or 139 amino acids, one of which was a blood isolate.

**Figure 5:**
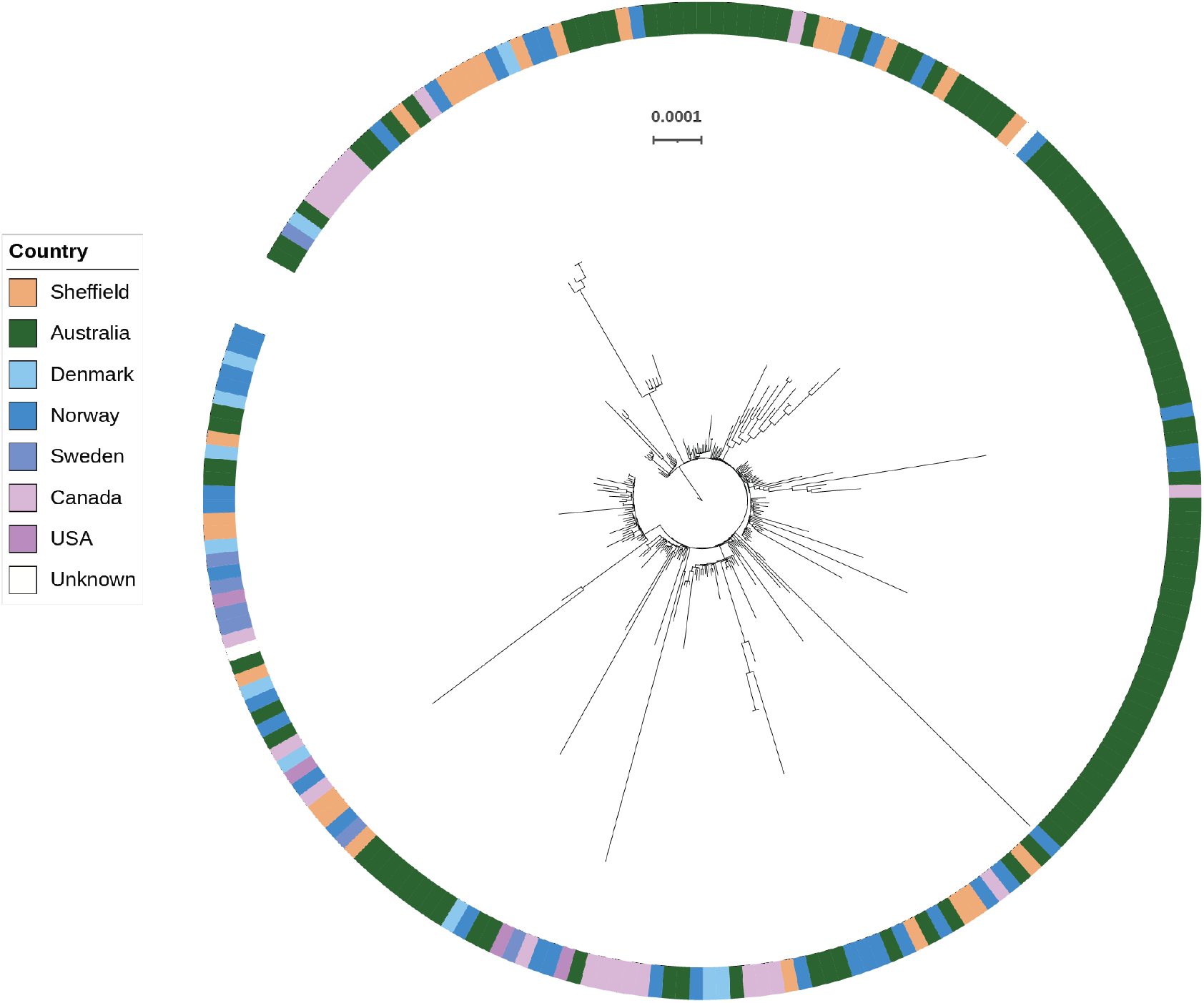
A phylogenetic tree of genome cluster 1 isolates. A maximum likelihood phylogenetic tree was constructed using RAxML (26) from core genome SNPs after mapping to the reference genome SDE096. Short read sequence data was obtained from NCBI SRA using the list of international isolates that belong to this lineage defined by Xie et al. 2025 (11). The ring indicates the country the isolates were collected from: Australia (n=115)(11), France (n=6)(17), Norway (n=74)(12, 18, 46), Denmark (n=11)(47), Canada (n=21)(14), USA (n=4)(48), Sweden (n=7)(49) and Unknown (n=2) (Rostock University Medical Centre, PRJEB34961), and also includes all 25 Sheffield isolates. The scale bar represents substitutions per site.

The SDE096 isolate was determined to carry the P601L variation in *pbp2X* conferring reduced penicillin sensitivity. This isolate also carried *ermA* and *tetM* which were found within a region between the conserved chromosomal genes *metQ* (D-methionine-binding lipoprotein, ACXAHW_06730) and *rlmD* (putative RNA methyltransferase, ACXAHW_07415), containing genes associated with plasmids and transposons (Figure 6) as well as a predicted incomplete phage. This incomplete phage region, along with the *tetM* and surrounding genes were also found in the published *S. pyogenes emm*230 genome (50) (Figure 6). Analysis of this region between *metQ* and *rlmD* in other isolates determined it to be highly variable and frequently associated with resistance genes (Supplementary Figure 3). Of the 37 for which we could extract this region over a single contig, three had *ermA* and *tetM* genes within this region, three had *tetM* alone and three had *ermA* alone. Within the GC-1 (*stG*62647/ST20) lineage, only four isolates carried any resistance genes at all, although two were from the same patient, but all carried them within this region (Supplementary Figure 4).

**Figure 6.**
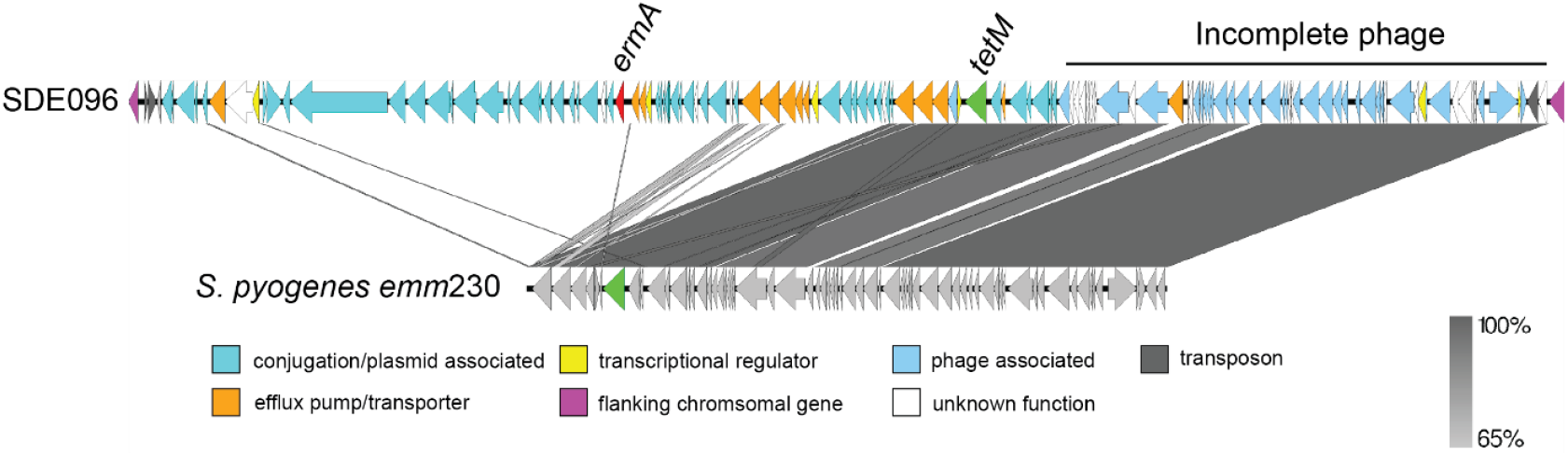
Schematic representation of the *ermA* and *tetM* region in SDE096. The *ermA* (red, ACXAHW_06910) and *tetM* (green, ACXAHW_07100) genes of SDE096 were found within a region located between the conserved chromosomal genes SDE096 ACXAHW_06730 (*metQ*) and ACXAHW_07415 (*rlmD*). The region downstream of *tetM* contained a predicted incomplete phage genome which shared homology with a region in *S. pyogenes emm*230 (NZ_CP035451.1). Figure generated using EasyFig. Percentage homology is indicated on the right.

### Diversity of the genomic cluster 2 isolates

The GC-2 lineage, predominantly consisting of ST17 isolates, with one single locus variant ST627 and ST282, and all Lancefield group G, was the most common cause of infections (n=34/96 isolates, 35.4%, counting three isolates from patient 1 and two isolates from patient 6 as one each). Eight different *emm*-types were found in this cluster, along with one non-typable due to a deletion within the *emm* locus. Gubbins analysis for recombination identified 692 genes across the genome that showed evidence of recombination from a total of 2,958 genes and a total of 275 blocks of recombination (Figure 7). As expected, given the diversity of *emm* genes in this cluster, *emm* was one of the genes. There was also a greater proportion of accessory genes in this cluster compared to GC-1 (Figure 1)

**Figure 7:**
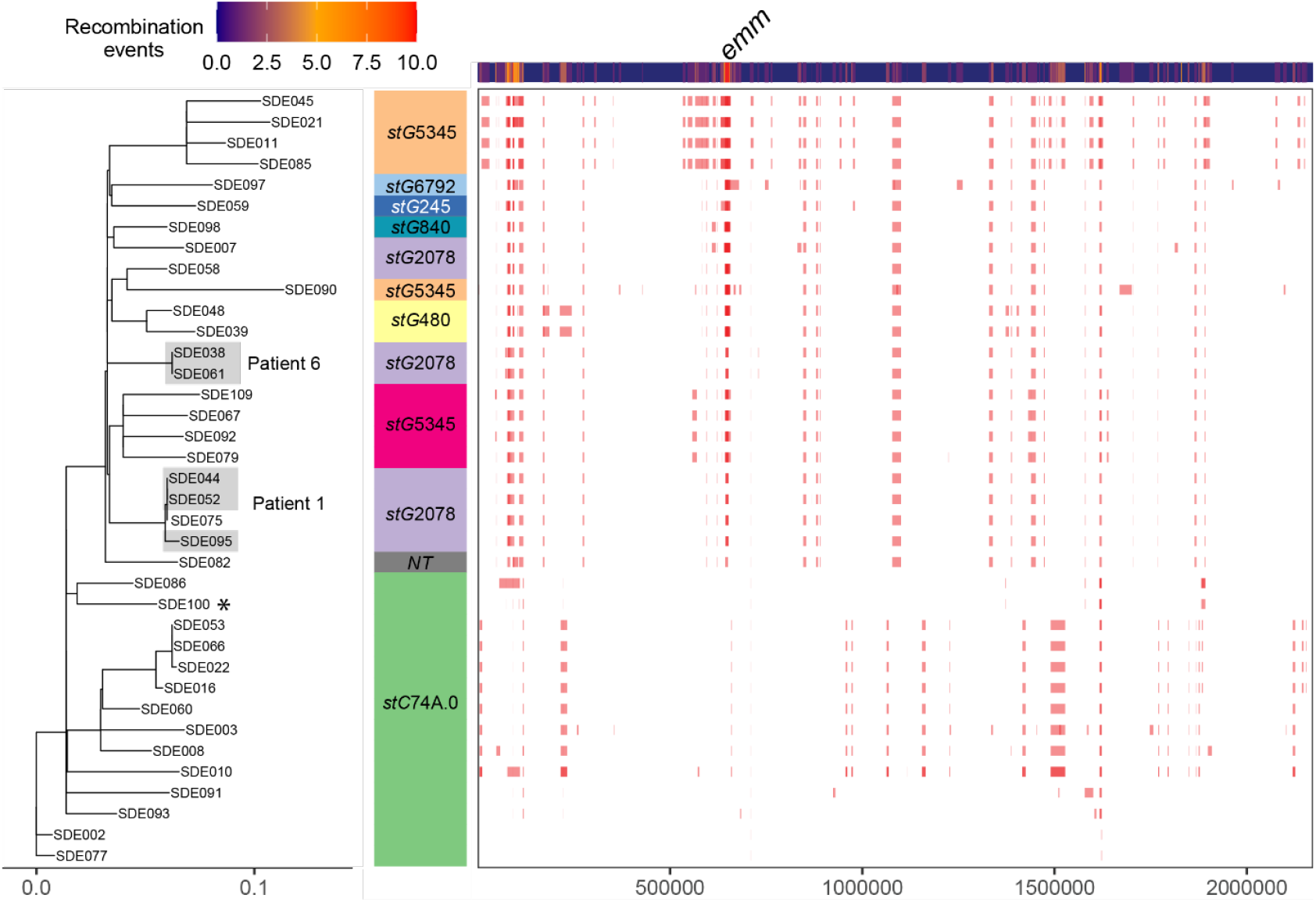
Regions of recombination in the SDSE genome cluster 2. Regions of recombination were determined using Gubbins with the *de novo* assembly for SDE100 as a reference (*). The tree on the left is a maximum likelihood tree generated from the Gubbins output (excluding recombination), the *emm-*type is given and on the right side are blocks showing recombination events with a heatmap of events above. The *emm*-type is given in coloured blocks. Three isolates from Patient 1 and two isolates from Patient 6 are shaded. SDE075 was not from Patient 1 but was closely related to the isolates from this patient.

All isolates for which we could determine the primary FCT type (26/34) all were FCT-A.II (26/30) similar to *S. pyogenes* FCT-6 (1, 40) with two fibronectin binding genes, *gfbA* and *fbpZ*, and the *pilB, pilA* and *pilL* genes. All but one isolate also had a secondary FCT region with another fibronectin binding protein gene and other pilin related genes. The *drsG/sicG* gene, which encodes for distantly related to Sic (DRS) and confers resistance to antimicrobial peptides, was found in 28/34 isolates within this cluster, and was not found in any isolates outside of this cluster (Figure 2).

The prevalence of antibiotic resistance genes was higher in this lineage than for GC-1, with 17 (44.7%) of isolates carrying at least one gene. Fifteen isolates carried *ermA*, five of these also carried *tetM*, one isolate carried *ermB* in addition to *ermA* and *tetM*, one carried *tetM* alone and one carried *ermB* alone. Three isolates (SDE022, SDE066 and SDE053) carried two copies of the *ermA* gene and both were located in the same region as *tetM*, within the *metQ*-*rlmD* variable region (Supplementary Figure 5). This region was the same for these three isolates and the region surrounding one *ermA* was found in seven others and two more with some differences, but differed substantially from another that had both *ermA* and *tetM*.

### Intrapatient variation

For six patients we had two separate isolates, obtained between 3 to 74 days apart, and for an additional patient we had three isolates, taken 4 and 36 days apart (Table 1). The three isolates from Patient 1 and the two isolates from Patient 6 were *stG*2078 and were present in GC-2 (Figure 7). Mapping to the *de novo* assembly of the earliest isolate for each patient identified that the isolates from Patient 6 (12 days apart) were identical, as were the first two isolates from Patient 1 (3 days apart), however the third isolate from Patient 1 (36 days later) differed by 5 SNPs. For all isolates within GC-2, excluding regions of recombination (Figure 7), there was a median and mean of 81 SNPs (range 0-144) between isolates, indicating that, as there was a low level of variation in the later Patient 1 isolate compared to the earlier ones, it was likely to be persistence of the same isolate rather than reinfection with a different isolate. However, SDE075 from another patient was identical to the early Patient 1 isolates, and so could have been part of a transmission event, however we do not have any epidemiological information to determine this.

**Table 1.** Genomic differences between isolates from the same patient.

| Patient | stG/GC | Days<br>apart | Changes | Impact |
| --- | --- | --- | --- | --- |
| Patient 1 | stG2078/<br>GC-2 | 3 | None | None |
|  |  | 36 | 5 SNPs | Y586C in BtuD (vitamin B12 import ATP-binding protein)<br>E150A in a prophage tail protein<br>E140G in ResA (TlpA disulfide reductase family protein) |
| Patient 2 | stG10/<br>GC-4 | 22 | 2 SNPs | R166H in SagC (part of Streptolysin S production) |
| Patient 3 | stG62647/<br>GC-1 | 57 | None | None |
| Patient 4 | stG652/<br>GC-7 | 74 | None | None |
| Patient 5 | stG62647/<br>GC-1 | 3 | None | None |
| Patient 6 | stG2078/<br>GC-2 | 12 | None | None |
| Patient 7 | stG652/<br>GC-7 | 28 | 2 SNPs | C123R in MntR (HTH transcriptional regulator) |

The isolates from Patient 3 and Patient 5 were *stG*62647 in GC-1 (Supplementary Figure 2) but both pairs were identical to each other, despite the 57 days between the Patient 3 isolates. The isolates from Patients 4 and 7 were from another cluster, *stG*652 but different ST (Figure 2), and although the pair from Patient 4 were identical despite the 74 day separation, the two isolates from Patient 7, 28 days apart, were two SNPs different, one non-synonymous (C123R) in *mntR* (a HTH transcriptional regulator) (Table 1).

### *S. canis* isolates

The five S. *canis* isolates we identified were Lancefield group G and all five were of a different ST, suggesting these were independent infections. One isolate carried two antimicrobial resistance genes (*ermA* and *tetO*), and another carried *tetO* alone. Consistent with the different STs, core genome comparison indicated a large genetic distance between the isolates with mean SNP distance of 8,063 (range; 57-80,625).

The penicillin binding proteins were also extracted from the *S. canis* assemblies and regions of variation identified. PBP1A had only two loci with variations A694T and G709S (Supplementary Dataset 1). PBP1B had 7 loci with variable amino acids N272K, V409I, A419T, V430V, S697P, A702T and N703P (Supplementary Dataset 1). There were three loci P68A, D127N and N764K in PBP2A that had variable amino acids all of which only found in one isolate compared to the other four. Finally, PBP2X had three loci that differ at position A108T, A281T and T535I (Supplementary Dataset 1). MIC testing of these isolates, however, did not indicate decreased sensitivity to penicillin.

## Discussion

In this study we investigated the genomic characteristics of SDSE isolates collected from all infection types in Sheffield, UK as no genomic data on SDSE isolates from the UK has yet been published. We found that, similar to other countries in Europe, North America and Australia, the GC-1 *stg*62647/ST20 lineage was common and there was a high level of diversity within other lineages.

Predominantly our isolates came from skin infections, followed by genital swabs, with a relatively low level of throat swab isolates. However, our collection period was June to October 2020 which was immediately following the COVID-19 pandemic lockdown period in the UK and therefore transmission patterns may have been different and throat swabs taken for microbiological identification of bacterial infections are likely to have been reduced. However, the numbers of bacteraemia cases notified to UKHSA for Lancefield group C and G during 2020 were no different to previous and post years (8). This is in contrast to group A streptococci where notifications of bacteraemia were substantially lower during 2020 compared to previous years (8). This was also the case for other countries as well (11, 13, 51, 52). We do, however, recognise that this collection period is likely to be different to a time without an active global viral pandemic.

Our SDSE population was diverse with 15 different genomic clusters, 20 different *emm*-types and 27 sequence types (STs). Like previous studies (5, 50) we found that *emm*-types and STs did not describe the population structure in the same way as they do for *S. pyogenes*. The most common genomic cluster (GC-2) comprised 8 *emm*-types and had a high level of recombination with 275 recombination events with an average length of ∼128 kbp. This was higher than GC-1, with 50 recombination events, and higher than that observed for major lineages of *S. pyogenes* (50, 53).

GC-1 (*stG*62647/ST20) was first reported in Argentina (16) but described in detail using whole genome sequencing by Oppegaard et al. in Norway (18). Although commonly reported as causing invasive disease in patients (3, 6, 10, 12, 18) other studies have shown no association with more severe outcome in patients over other lineages (11, 14, 54) and varying mortality in mouse models of infection (17). Our *stG*62647/ST20 isolates were part of the international lineage genetically predicted to have emerged in 1956 before rapidly increasing in the population (11). Characteristically of this lineage, all our isolates carried the disrupted *silB* which may influence virulence, although there are many other mechanisms influencing virulence in this lineage (55). In our sampling period we had very few isolates from blood infections (n=7) and none of these were GC-1 isolates, however it is not known what the prevalence of this lineage is on a national scale and over a much wider time period.

In addition to the disruption of *silB* in GC-1 isolates, we also identified three isolates from other genomic clusters that had premature stop codons in this gene, potentially influencing virulence of these isolates. Mutations in the two-component regulatory system CovR/S also have the potential to influence virulence and have been associated with severe disease and STSS (42-44, 56). We identified six amino acid variations in CovR (5.7% of isolates), but 18.3% of isolates had amino acid variations in CovS. It is not clear what the impact of the amino acid variations in CovR or CovS may be as, like we have observed for *S. pyogenes*, they could be associated with lineages and have no effect on function (57-59).

Tetracycline resistance genes (*tetM/tetO/tetW*) were found in 36.5% (n=38) of isolates, which is a similar proportion to what we previously found in *S. pyogenes* from skin infections in Sheffield (37.3%) (57). Other reports of tetracycline resistance in SDSE have been between 28% and 62% (6, 13, 15, 16, 54, 60). The proportion of our SDSE isolates carrying *ermA* or *ermB* potentially conferring macrolide resistance was also high (35.6%) and ∼5 times higher than in the Sheffield *S. pyogenes* population (6.5%) (57). Similar levels have also been found in other studies (6, 16, 54). Whilst group A streptococci (*S. pyogenes*) are now included on the WHO priority pathogen list because of the growing concern of macrolide resistance (61) there should also be a concern for macrolide resistance in group G/C streptococci as well.

A high proportion of the *tet* and *erm* genes were present in a variable region between the conserved chromosomal genes *metQ* and *rmlD*. In SDE096 and other isolates there were several genes in this region predicted to function as conjugal transfer and mobilisation proteins, indicative of a plasmid or extrachromosomal element. A previous study did find an extrachromosomal circular element in an *stG*62647 strain that had an attachment site homologous to *rmlD*, however it could not occupy that region as there was already a different mobile genetic element in that site (17). It is not clear if this region in any of our isolates could mobilise and replicate extrachromosomally or if there are additional extrachromosomal elements. For 37/104 of our isolates the region assembled over a single contig but for other isolates it would require long read sequencing to accurately determine the composition of this region, as well as any extrachromosomal elements. Some isolates carried resistance genes outside of these regions and these were associated with highly variable ICEs that we did not describe in detail here. However, it does indicate that the acquisition of resistance genes by SDSE can be via many routes and with several other bacterial species acting as potential donors.

We found a number of variations within the penicillin binding proteins although only P601L in PBP2X showed a reduction in sensitivity to penicillin, as had previously been found with *S. pyogenes* (37). The MIC for the two isolates with the P601L variation was the highest at 0.03 mg/L, which is the breakpoint as defined by EUCAST (https://www.eucast.org/). It is possible that, although this variation in itself does not confer high level resistance nor has reduced sensitivity to a level that might have clinical impact, it is possible that these mutations are the starting point for such resistance to develop. These two isolates were obtained from the same patient, 57 days apart and had remained unchanged genetically. Whether the presence of the P601L mutation impacted on the lack of infection clearance in this patient is unclear as we do not have the clinical information available to know what treatment was given and for how long. We do not know the history of this patient and therefore can only hypothesise that the variation in PBP2X arose during infection although, as no other isolates were identified carrying this variation, it seems likely that this was the case.

The lack of genetic change from 4/7 pairs of isolates from the same patient indicated long term persistence within the host. The earliest isolate from Patient 4 was from blood culture and this was followed by a skin swab isolate 74 days later, yet these were identical, indicating long-term persistence within a patient even with the potential to cause invasive disease. When genetic changes did occur, it did not seem to be related to length of time or genomic cluster, indicating a host factor. It is not clear what the impact of any of the genetic changes observed would be to the isolate but there was no evidence for changes that would alter antimicrobial sensitivity. Although it is possible that the variation in *mntR* that arose in Patient 7 could alter metal transport which in turn could alter resistance to oxidative stress (62). As no other clinical information can be obtained for any of these isolates, it is not known if there were other infections prior or post to those we collected, nor do we know what kind of therapy the patients underwent and whether there was any evidence of treatment failure. However, long-term persistence and the potential for repeated treatment failure is a concern.

An additional public health concern was the finding that five isolates were *S. canis*, a pathogen thought to rarely infect humans. Originally identified in dogs, *S. canis* can infect a number of different animals, including cats, and therefore household pets may be a source of transmission to humans (33). Prevalence in humans may be underestimated as *S. canis* is often diagnosed as a Lancefield group G *Streptococcus* rather than confirmed at the species level, as was the case in our study prior to genome sequencing. Given the fairly high proportion (4.5%) we found, coupled with the fact that two isolates carried tetracycline resistance genes and one of these also carried a macrolide resistance gene, it is important to note that *S. canis* can cause infections in humans and should be highlighted as a public health concern. It also demonstrates the importance of whole genome sequencing in routine diagnostics for correct identification and speciation of closely related pathogens.

Similar to other studies (1, 39, 40), we found two FCT regions in the SDSE isolates and, although we could not determine the exact FCT type for all of them due to contig breaks in the genome assembly, a primary FCT was found in all isolates while a secondary FCT was only found in 82.5% of isolates. For *S. pyogenes* there have been links made between FCT and tissue tropism (38) but there is limited data on SDSE, therefore it is unclear what these two FCT regions contribute to tissue tropism. Studies have also shown that the architecture of the primary FCT could be predictive of SDSE adhesion properties but its role in tissue tropism was not clear (40). Both FCT regions contained genes encoding proteins for pilus formation, as is found in *S. pyogenes*, but it is unknown whether they would produce two separate pili, one from each of the FCT regions.

Overall, we have identified that, like other countries, genome clusters GC-1 and GC-2 are highly prevalent, accounting for 60.6% of all the isolates. The high level of diversity within the SDSE population makes it difficult to monitor with typing methods, like *emm* or MLST, alone and whole genome sequencing is required to fully understand the population. It is still unclear as to why GC-1 and GC-2 are so common in the population and what bacterial factors might be driving this. Genome sequencing of SDSE on a national scale is needed to better understand the UK population, in particular to determine any invasive disease potential associated with GC-1 or GC-2. It would also be of interest to confirm the low level of throat infections caused by SDSE in comparison to skin infections and if this differs to *S. pyogenes* for which we have found more throat infections compared to skin infections (30).

## Author contributions (CRediT)

**SYB:** data curation, formal analysis, investigation, software, visualisation, writing – original draft, **HK:** resources, methodology, investigation. **SJ:** investigation. **RC:** software, investigation. **LRG:** investigation. **LT**: resources and methodology, **DP:** resources. **TIdS:** resources, funding acquisition, writing – review & editing. **CET:** conceptualisation, data curation, formal analysis, funding acquisition, investigation, methodology, project administration, supervision, visualisation, writing – original draft, writing – review & editing.

