## Supplementary Figures for "Molecular analysis of Lancefield group C/G streptococci causing human infections in Sheffield, UK"

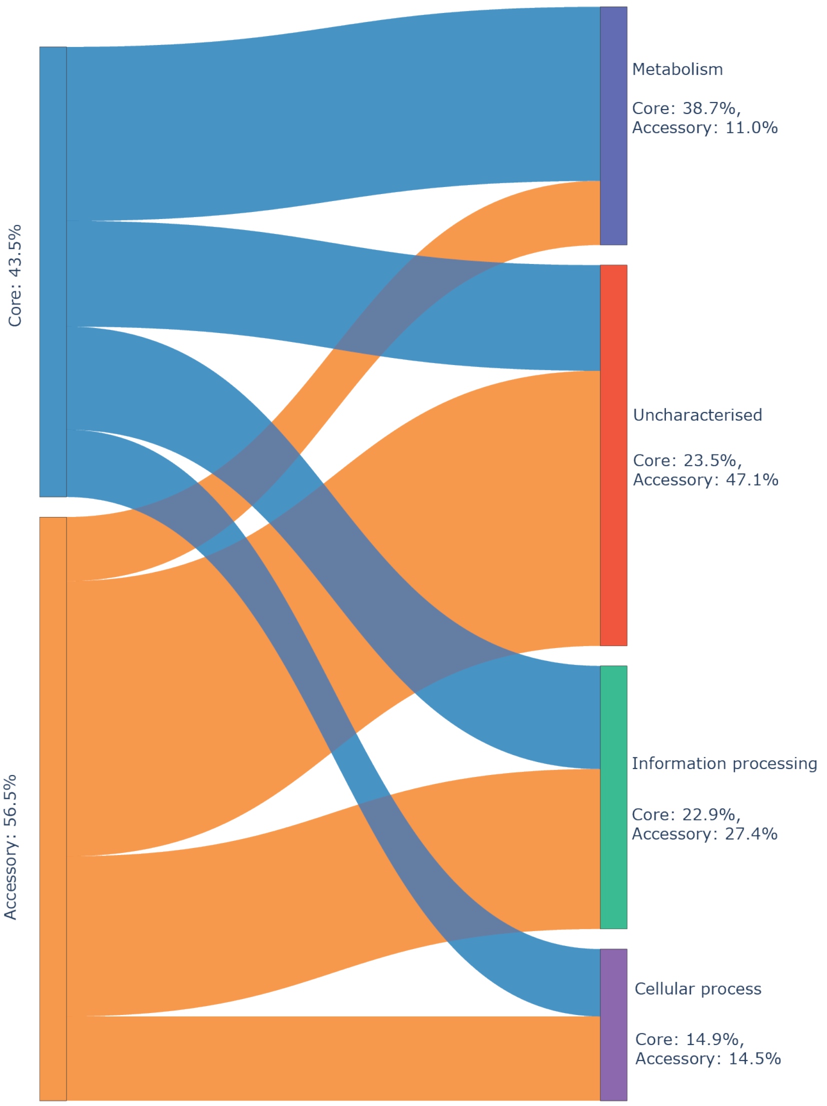


**Supplementary Figure 1. Predicted functional characterisation of the core and accessory genome of the SDSE isolates.**

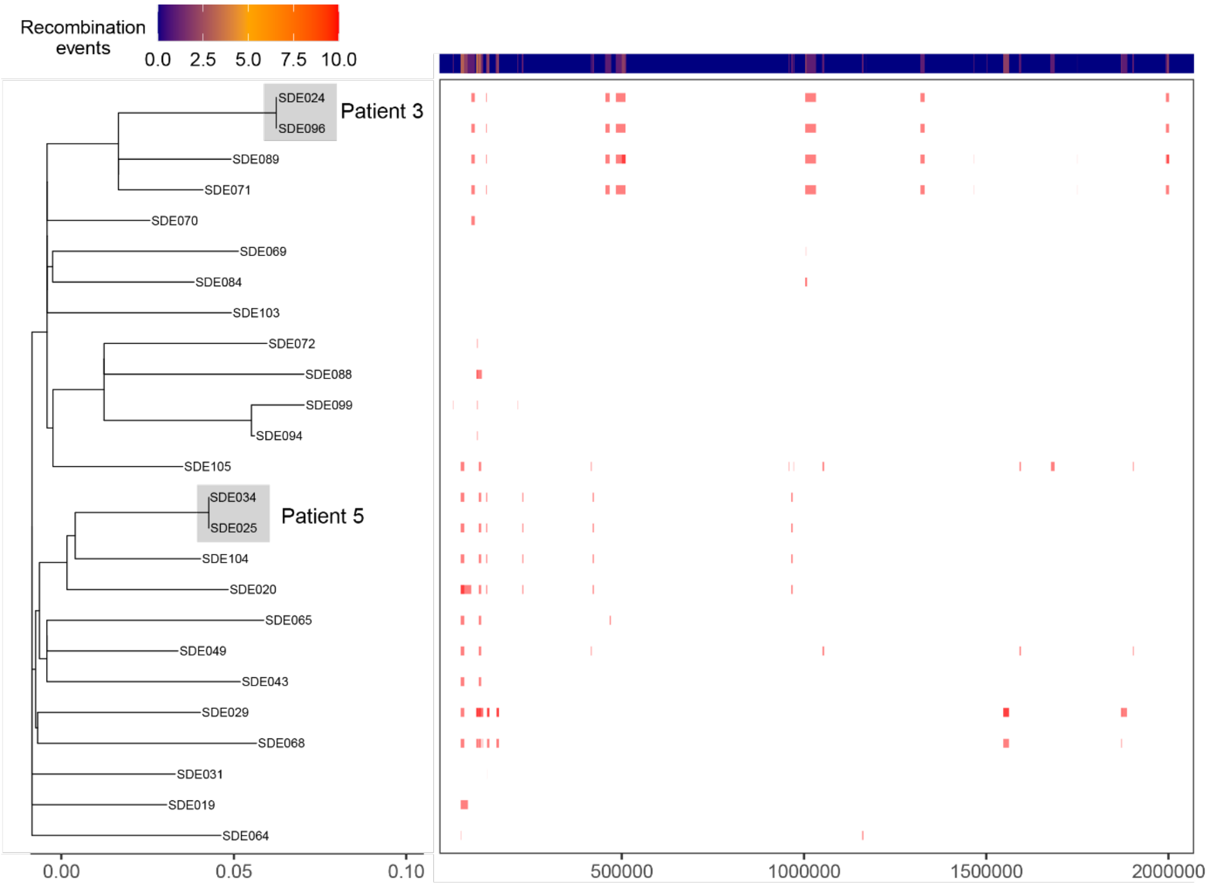


**Supplementary Figure 2. Recombination within GC-1.** Gubbins analysis for recombination within the GC-1 lineage using SDE096 as a reference. Two pairs of isolates came from Patient 3 and Patient 5 (highlighted in grey).

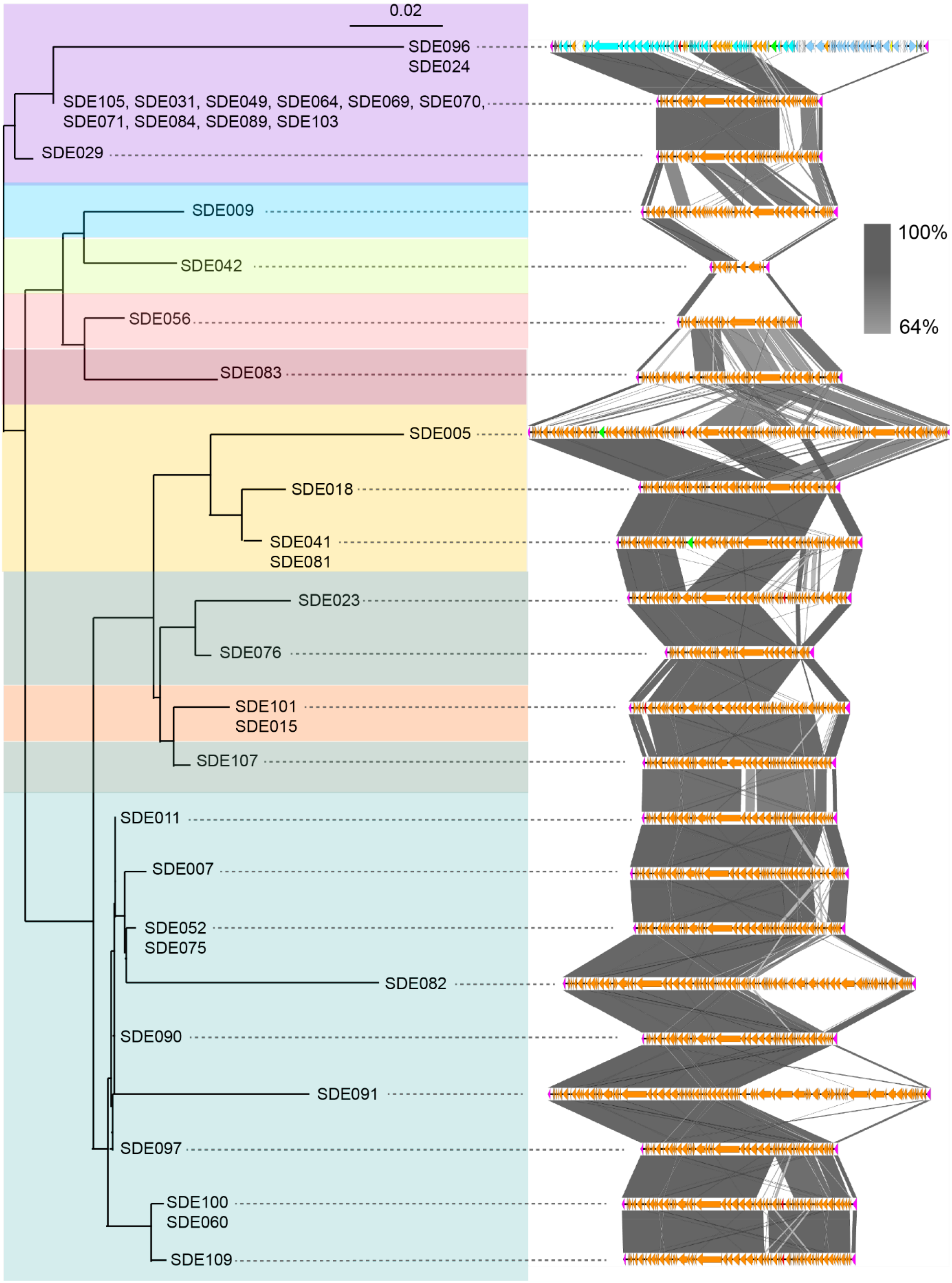


**Supplementary Figure 3. Diversity of the *metQ* to *rlmD* variable region.** The region between *metQ* and *rlmD* was extracted from 37 isolates where it was assebmled within a single contig. A phylogenetic tree was generated using gene presence/absence within the *metQ-rlmD* region. This region was quite variable even within genomic clusters (represented by branch shading: each colour represents a genomic cluster). With the exception of SDE096, only *tetM* (green), *ermA* (red) and flanking *metQ* and *rlmD* (pink) genes are indicated. Percentage homology indicated. The top cluster (SDE096, SDE015 and SDE029) are GC-1 *stG*62647/ST20 and the bottom cluster (SDE011-SDE109) are GC-2.

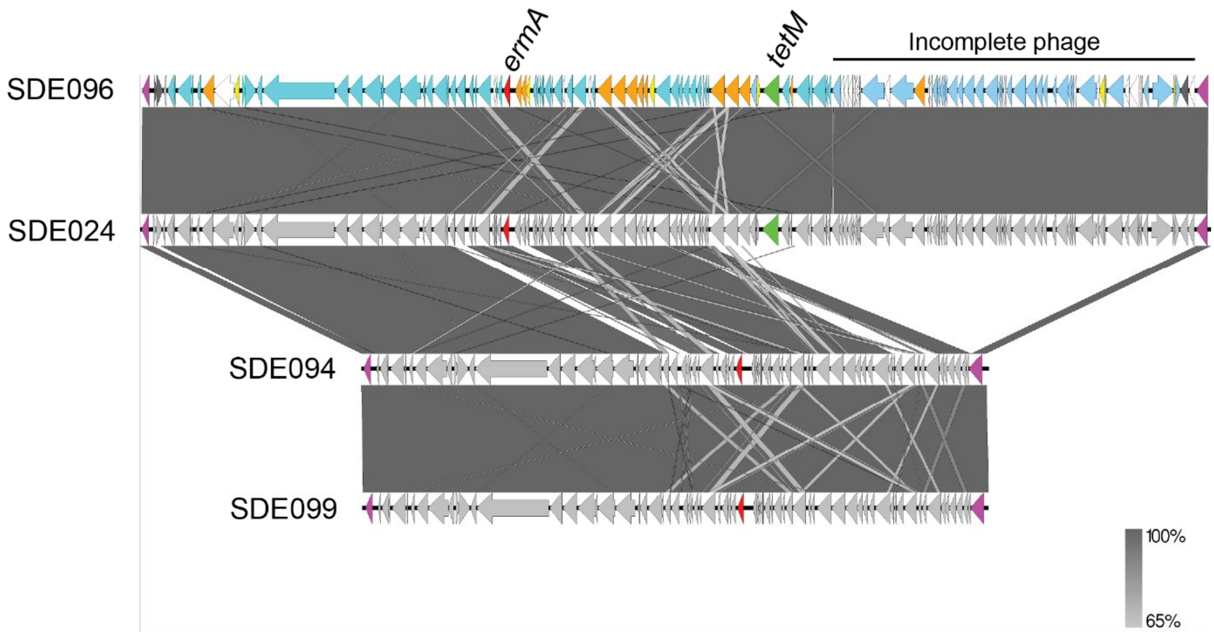


**Supplementary Figure 4. Genomic cluster 1 *metQ-rlmD* region.** Only four isolates within this lineage had resistance genes, two from the same patient (SDE096 and SDE024), and they were all present within the region of variation between the chromosomal genes *metQ* and *rlmD* (pink). This region was almost identical between all four isolates with the addition of the prophage-like region in SDE096 and SDE024 with the *tetM* gene (described in Figure 6). Apart from SDE096, only the flanking genes (pink), the *ermA* gene (red) and the *tetM* gene (green) are coloured by function. This region in SDE094 and SDE099 was split over two contigs, reordered against SDE096, with the contig break immediately after *metQ*, therefore we cannot rule out the possibility of additional genes within this region. Figure created using EasyFig.

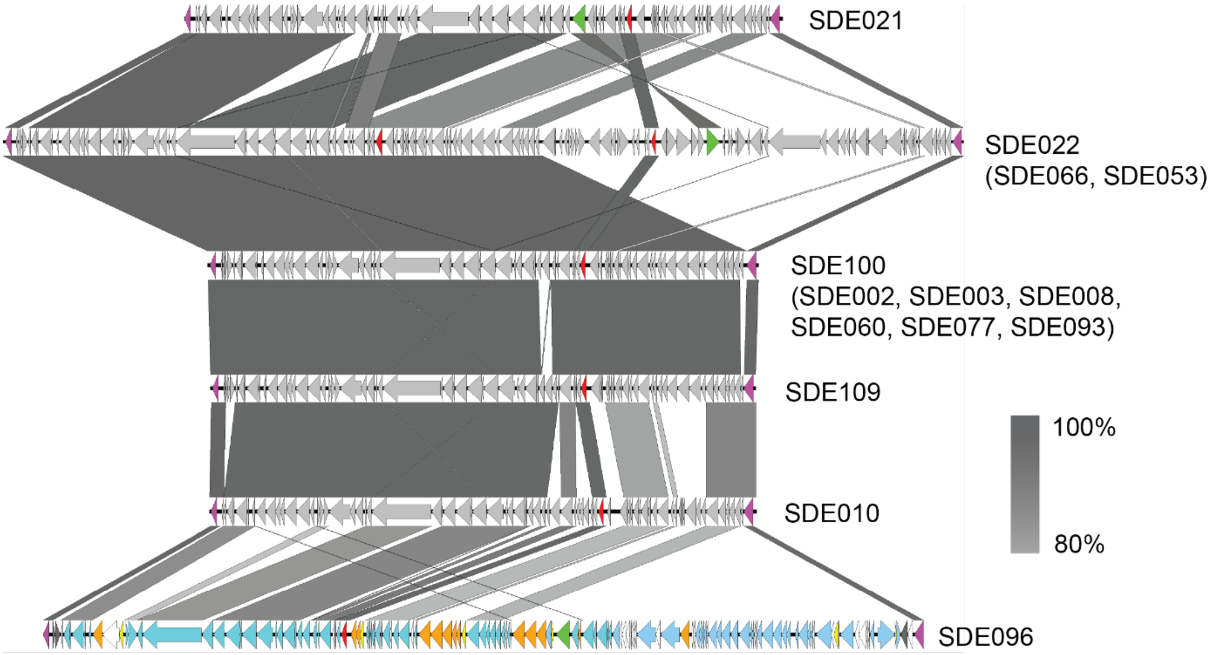


**Supplementary Figure 5. Genomic cluster 2 *metQ-rlmD* region.** A comparison was made of the region between *metQ* and *rlmD* for those GC-2 isolates that carried *ermA* or *tetM*. With the exceptions of SDE060, SDE100 and SDE109, this region was split over several contigs which were then ordered against the SDE096 complete genome. Therefore this is a predicted structure and we cannot rule out the possibility that there could be other genes present in these regions or that this prediction of this region is incorrect. Apart from SDE096, only the flanking genes (pink), the *ermA* gene (red) and the *tetM* gene (green) are coloured by function. SDE022 and SDE100 are representing the same region in other isolates listed in brackets. Figure generated using EasyFig.
